# Analysis of follow-up data in large biobank cohorts: a review of methodology

**DOI:** 10.1101/2023.06.14.542699

**Authors:** Anastassia Kolde, Merli Koitmäe, Meelis Käärik, Märt Möls, Krista Fischer

## Abstract

This study focuses on key methodological challenges in genome-wide association studies (GWAS) of biobank data with time-to-event outcomes, analyzed using the Cox proportional hazards (CPH) model. We address four primary issues: left-truncation of the data, computational inefficiency of standard model-fitting algorithms, relatedness among individuals, and model misspecification. To manage left-truncation, the common practice is to use age as the timescale, with individuals entering the risk set at their age of recruitment. We assess how this choice of timescale influences bias and statistical power, under realistic GWAS conditions of varying effect sizes and censoring rates. In addition, to alleviate the computational burden typical in large-scale data, we propose and evaluate a two-step martingale residual (MR) approach for high-dimensional CPH modeling. Our results show that the timescale choice has minimal effect on accuracy for small hazard ratios, though using birth age as the timescale-ignoring recruitment age-yields the highest power for association detection. We find that relatedness, when ignored, does not substantially bias effect size estimates, while omitting key covariates introduces significant bias. The two-step MR approach proves to be computationally efficient, retaining power for detecting small effect sizes, making it suitable for large-scale association studies. However, when precise effect size estimates are critical, particularly for moderate or larger effect sizes, we recommend recalculating these estimates using the conventional CPH model, with careful attention to left-truncation and relatedness. These conclusions are drawn from simulations and illustrated with data from the Estonian Biobank cohort.

## 1 Introduction

As time goes on, the data volume in large-scale population-based biobanks is increasing exponentially. Although the recent decades have seen tremendous increases in sample size, a similarly valuable data expansion results from prolonged follow-up time and the ability to link the -omics databases with incident disease data from electronic health records. Therefore, a large proportion of Genome-Wide Association Studies (GWAS) are mainly focused on discovery of genetic variants associated with the risk of incident diseases. For that purpose, one needs to apply regression modelling methodology that is designated for censored time to event data, rather than using simple methods like linear or logistic regression models [1, 2, 3]. Here, the Cox Proportional Hazards (CPH) modelling [4] has become a standard in biomedical fields due to its robustness to distributional assumptions and interpretation of parameter estimates in terms of Hazard Ratios (HR-s).

Despite its robustness, CPH modelling is not assumption-free. Therefore there is a need for a review of applicability of this method in the context of large population-based biobanks. Our aim is to identify sources of possible biases as well as realistic magnitudes of them in typical GWAS settings.

The most discussed assumption in the context of CPH model is the proportional hazards assumption, stating that the multiplicative effect of a risk factor on the hazard of the outcome event is staying the same throughout the scale of the follow-up time. Recently it has been pointed out that this is rarely true in practice – on the contrary, the hazard ratios are almost always time-varying. Therefore the HR from a CPH model should be interpreted as a weighted average of the true HRs over the follow-up period [5, 6]. This could be easily acceptable in GWAS, unless there is a reason to believe that some genetic variants have a drastically different effect on the risk during different segments of the follow-up time.

Some other, often ignored assumptions are related to the special features of the biobank data. First, we note that the time of recruitment is usually not a relevant baseline timepoint regarding the outcome event (unlike in clinical trials, where follow-up often starts at diagnosis). As the genomic data stays largely constant throughout the lifetime of an individual, date of birth may seem as a logical time origin for a GWAS. Using age at the outcome event as the outcome variable can, however, lead to another problem called left-truncation or immortal time bias [7, 8], as the analysis is still conducted conditionally on the fact that the individual was alive at the time of recruitment and free of diseases (sometimes including the outcome event) that would have prevented the recruitment. To properly account for left-truncation, one should use methods that use age as timescale, but consider the individual as being at risk only during the time from recruitment until the outcome event (or end of follow-up)[7].

A common feature of biobank cohorts is genetic relatedness of the participants, violating the assumption of independent observations in the sample. For continuous trait GWAS, the use of mixed linear models has been recommended in such cases [9, 10, 11, 12, 13]. Similar approach could be used in survival analysis (mixed effects Cox regression, frailty models) [14].

When the outcome event is an incident disease, mortality due to other causes will always be a competing event – censoring the individuals where the follow-up ended due to death, ignores the assumption of independent censoring. In this case, one may consider using a proper model for competing risks (Fine and Gray model). However, when the focus is not on risk prediction, but on parameter estimation, censoring the competing outcomes is still acceptable [15, 16].

As the biobank cohorts are mostly not random samples from the population, also other sources of selection bias are likely [17], that could sometimes be addressed by proper use of sampling weights.

In addition to the biases resulting from sample design, also some computational approaches used in GWAS may become sources of bias. Due to the significant increase in the size of genotyped samples over the past few decades, both in terms of the number of genotyped subjects and the number of genetic variants genotyped or imputed, most of proposed tools for running CPH modeling in GWAS setting have become computationally prohibitive and not easily scalable[18, 19, 20, 21]. In some studies, a two-stage approach involving martingale residuals that dramatically reduces the computing time, has been used [22, 23]. Although it has shown to perform well in some simulations, it is still unclear, whether it results biases in parameter estimates in some realistic cases.

In classical linear regression with independent covariates, consistent and unbiased estimates for the remaining coefficients can be obtained even if some covariates are omitted. However, this property does not extend to non-linear models, including logistic regression and CPH model neither in randomized or observational setting [24, 25, 26, 27, 28, 29].

In a CPH model, omitting an important risk factor for the outcome leads to violation of the proportionality assumption with respect to other variables in the model, leading to omitted variable bias, which can significantly distort estimates and conclusions.

In GWAS context one still needs to keep in mind the main task of identifying the potential disease-associated variants in the set of a large number (often more than 20 millions) of genotyped variants. As the focus is on hypothesis testing rather than precise effect estimation, small biases are not a cause of concern, if the nominal alpha-level is still retained after accounting for multiple testing. Therefore, if there is a trade-off between bias and power, a biased estimator may be preferable if it leads to greater power.

The main aim of the present study is to assess the magnitude of bias and power in realistic GWAS settings, where the “naive” CPH model is used, while ignoring left-truncation and/or relatedness of individuals, possibly using the martingale residual approach to speed up the computation. We explore these questions analytically and also by a simulation study, clarifying the need for various bias-reduction measures in GWAS settings.

Finally we also address the option to combine participant genotypes and parental outcome data, when the biobank cohort is relatively recent and the number of events still low (especially for mortality outcomes). Clearly, the estimated HRs in this case will not be unbiased. We derive the expression for the bias analytically and demonstrate the performace of this approach in a small-scale simulation.

## 2 Sources of bias

### 2.1 Timescale choice

#### Bias due to left-truncation

Here and hereafter we are considering the Cox proportional hazards (CPH) model defined as

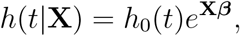

where *h*_0_(*t*) is the baseline hazard function at time *t*, **X** is the matrix of covariates and ***β*** is the vector of the parameters.

To estimate the parameter ***ψ*** = *e*^***β***^ using the CPH model, one needs to find the value of ***β*** that maximizes the partial likelihood function:

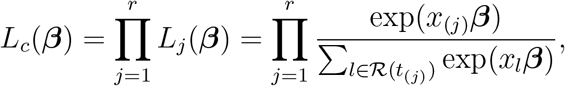

where *x*_(*j*)_ is the value of *X* for an individual who died at the *j*th observed death time *t*_(*j*)_, *x*_*l*_ are the values of *X* for the individuals in the risk set at *t*_(*j*)_ and *r* is the number of events. Note that the risk set ℛ (*t*_(*j*)_) consists of individuals observed to survive up to *t*_(*j*)_. Obviously, for unbiased estimation the risk set in the cohort should be a random sample from the corresponding risk set in the population. However, when time of birth is used as the time origin rather than the time of recruitment, individuals who were recruited at age *t* + *δ* would be included in the risk set for an event at age *t* for any *δ* > 0. Thus the risk set at *t* is partly selected conditionally on events (recruitments) that occur after *t* and thus also conditionally on the fact that the individuals survive (and have no outcome events) before *t* + *δ*. As a result, low-risk individuals will be over-represented in that risk set (as they have more chance survive up to time *t* + *δ*), leading to overestimation of the hazard ratio.

Still, while using time of birth as time origin and ignoring left truncation, no bias is anticipated under the null hypothesis (***ψ*** = 1 or ***β*** = 0). Therefore the question remains, whether the approach of using age as timescale and time of birth as time origin could still be valid or even preferred for hypothesis testing due to potentially better power.

#### Potential bias with time since recruitment as timescale

A standard “textbook” approach would be to use time since recruitment as timescale, while the effect of age is accounted for by proper adjustment. Korn at al [8] have determined two sufficient conditions – when either one is satisfied, the age-adjusted CPH models with time since recruitment as timescale and CPH models with age as timescale give the same results:

1. The baseline hazard function *h*_0_(*t*) can be presented as an exponential function: *h*_0_(*t*) = *c* · exp(*γt*) for some *c*> 0 and *γ*.
2. The covariate of interest and age at recruitment are statistically independent.

If neither of the two conditions is fulfilled, Korn et al. suggest using CPH models with age as timescale, as they believe the outcomes to change more as a function of age rather than a function of time since recruitment.

The first condition holds when hazard of the outcome is expected to increase rapidly as a function of age, following a Gompertz distribution (often appropriate for human mortality data).

Korn et al. did not provide a formal proof for the second condition, whereas Thiebaut et al. [7] found that mismodeling of age as an adjustment factor in a follow-up-dependent CPH model rather than as a timescale could also result in bias, on the contrary to the reasoning by Korn et al. Other authors have shown that bias can be detected even when a variable independent from the variable of interest has been omitted from the CPH model [24, 25, 26, 27]. One can argue that while modelling all-cause mortality, the second condition basically states that the covariate of interest cannot affect mortality as the distribution of this covariate would otherwise change with age, making them dependant. The distribution stays invariable, when the covariate does not affect mortality.

Thiebaut et al. also suggest using age as timescale rather than time since recruitment, as the underlying mechanisms of these models are different. They point out that usually the time when subject comes under observation does not coincide with the time when the subject becomes at risk for the outcome of interest. This is especially true in the biobank context.

Again, our question is related to the practical implementation of these findings in the context of GWAS – what is the effect of timescale choice in a range of realistic settings on bias in parameter estimates as well as on power to detect an association. We will try to shed some light on these questions using a simulation study.

### 2.2 Dependent observations

Independence of observations is a central assumption in most modeling approaches, including the CPH model. However, in population-based biobanks it is common to encounter genetically related individuals, which violates this assumption and can introduce biases. A general approach to address population structure in GWAS is to add principal components (PCs) as covariates [30, 31]. While PCs primarily address population stratification, they also might help in identifying and accounting for cryptic relatedness within the sample. However, specialized methods are preferable for explicitly adjusting for related individuals alongside population stratification. These methods include using a kinship matrix [9], linear mixed models [10, 11, 12, 13], or frailty models [14], which are more effective than PCs alone, but are computationally prohibiting.

### 2.3 Omitting covariates

The CPH model relies on the fundamental assumption that the effect of each covariate is proportional over time and relative to other covariates in the model. Although covariate effects can be approximately proportional in reality, apparent non-proportionality often results from model misspecification [24]. Even in randomized trials where all known and unknown confounders are balanced between study arms, omitting a covariate can lead to bias in treatment effect estimates. This bias is particularly problematic in observational studies, where neither the observed covariates nor unmeasured confounders are balanced across groups with different exposure levels (or genotypes). Consequently, the risk of omitted variable bias is significant, potentially leading to asymptotic biases. This results in systematically incorrect estimates, which can mislead conclusions about the relationships between variables. The severity of bias depends on the distribution of the omitted covariate, strength of its effect and censoring. Although one cannot directly adjust for unmeasured covariates, their potential impact can be assessed by sensitivity analyses [29], but that is hardly ever done in GWAS setting.

### 2.4 Other issues

In addition to the problems mentioned above, there are various other potential sources of bias that may affect the final conclusions, depending on the research question – we list them here for completeness, but ignore in subsequent analysis.

When the outcome of interest is not death, but an incident condition, one should be aware of **competing risks**, such as death due to another cause. While treating competing events as censoring, however, the hazard ratios are still unbiased in general [16], but care should be taken in absolute risk estimation tasks.

The assumption of **proportionality of hazards** has been discussed above in the context of omitted covariates, but the risk factors (including genetic variables) themselves may also have a time- or age-varying effect on hazard. However, this is again an issue that becomes important in absolute risk prediction, while for variant discovery one may accept that an average effect over time is estimated.[5].

Recently it has been pointed out by several authors, that population-based biobanks are mostly non-random samples and therefore subject to **selection bias** [17, 32, 33, 34]. We agree that it is an important issue that should be taken into consideration in biobank-based studies, regardless of the type of variable (time to event or other) or method of analysis.

## 3 Ways to increase power and computational efficiency

### 3.1 Parent-offspring data

Joshi et al. [22] have combined parent-offspring data to increase power of discovery. Biobank cohorts with short average follow-up time are underpowered for the analysis of participant lifespan data, due to the low number of outcome events (deaths). However, if family history at recruitment is collected and parental ages at death are known, they can be combined with subjects’ genotype information.

If the age span of recruited subjects is sufficiently wide, a large proportion of them is likely to have parents who are either relatively old or already deceased. Therefore the use of parental data leads to lower censoring rates and higher power for genotype effect detection. As each allele in a SNP is inherited by offspring with the probability of 50%, one can assume that using parental lifespan along with offspring genotype will result in estimates with magnitude of about half of the true effect size.

However, we have shown that the proportionality assumption in this case does not hold. We have derived equations for bias and show that the bias will increase, if the minor allele frequency and/or the effect size *β* increases. The derivation is explained in detail in the Supplement Section 1.

The approach of combining parental and offspring data has mainly been used for power gain, but our simulations showed that it is justified only when the parental event rate is at least 4 times higher than the event rate of the recruited subjects.

### 3.2 Two-stage modeling via martingale residuals

To overcome computational challenges and leverage GWAS tools tailored for continuous phenotypes [35, 36, 37, 38], we will examine performance of a two-step modelling approach proposed by Joshi et al. [22] for a biobank setting. The idea of the method is fairly simple – instead of running a CPH model for every SNP, a single CPH model encompassing all the non-genetic and technical covariates is fitted in the first step. For that model, martingale residuals (MR) are obtained (Supplement, Section 2.3.2). As pointed out by Therneau et al. [39], the association between MR and a covariate omitted from the linear predictor of the initial model yields estimates that align with the coefficients in the CPH model. Thus a test of a linear association between the MR and a genetic variant could potentially be used to detect an association between the variant and the outcome phenotype, reducing the association testing to a simple linear regression task.

## 4 Results of the simulation study

We will simulate different scenarios similar to real life biobank data in order to determine if and how the above mentioned methodological choices in the CPH model affect the results. We will study the bias and power under various minor allele frequencies (MAF), effect sizes and censoring rates. Timescales used:

- timescale **TB** – time since birth;
- timescale **TR+A** – time since recruitment, age-adjusted;
- timescale **TA** – age as timescale (accounting for left-truncation).

More detailed simulation strategy is given in Supplement Section 2.1.

Our main aim is to compare effect size and significance of a SNP using the conventional CPH model and two-step MR approach. The impact on the working range of the approach is examined. The study aims to determine the effects of censoring and MAF on the performance of the two-step MR approach.

We will compare the models performances by:

- bias - difference between real effect size and the estimated effect size;
- power - probability of detecting a significant effect, when it is present;
- coverage - probability that the true effect size lies in the confidence interval of the estimated effect size.

### 4.1 Effect of timescale choice on bias and power in CPH model

CPH with TA (Figure 1) is the only one that results in unbiased estimates for *β*_1_. In addition, this model exhibits the best coverage of the 95% confidence interval for true effect size.

**Figure 1.**
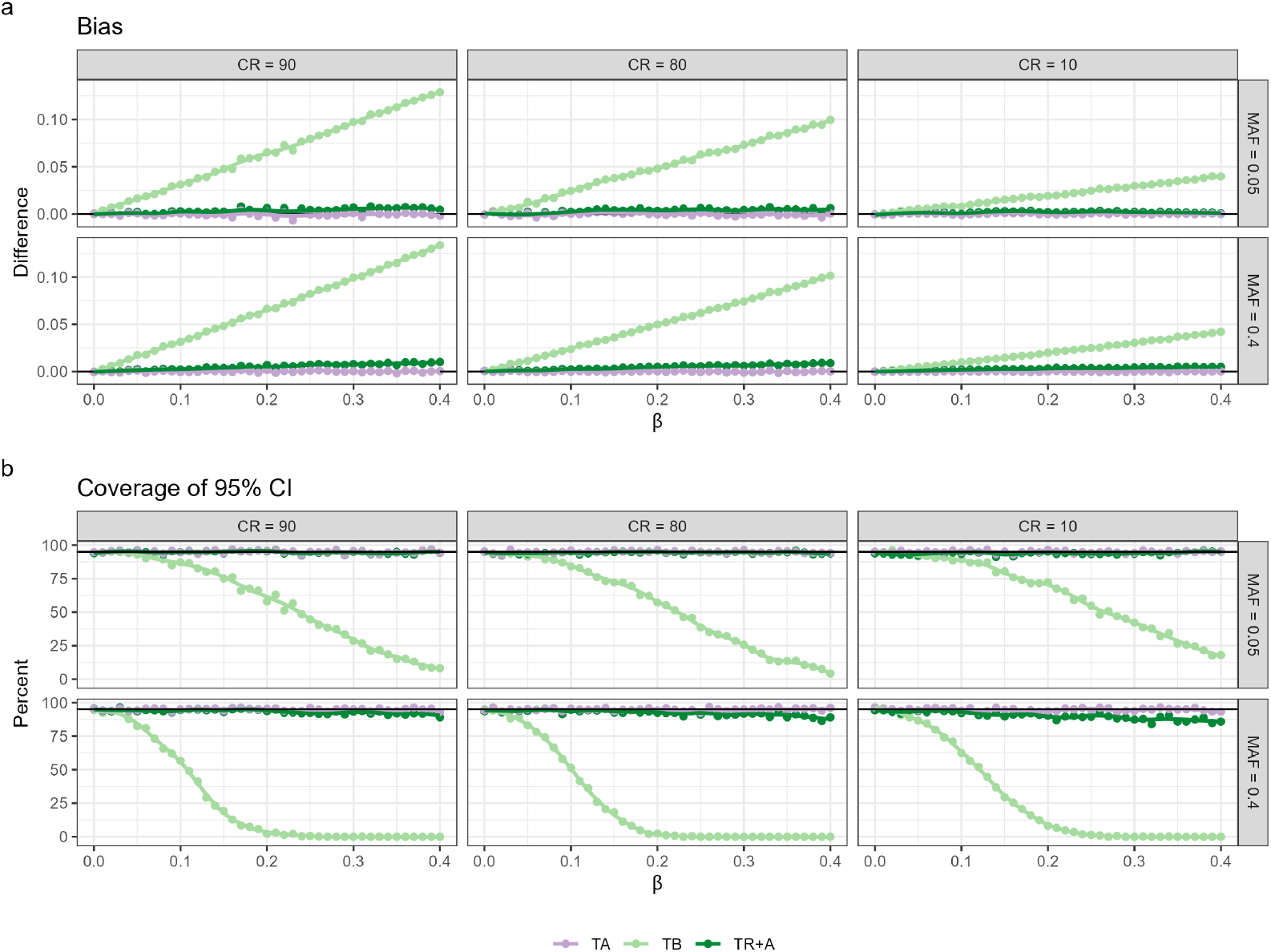
Performance metrics of CPH with different timescalesage, account for left-truncation (TA), time since birth (TB) and time since recruitment with age adjustment (TR+A) under different censoring rates (CR) and minor allele frequencies (MAF), encompassing the Bias and Coverage.

CPH with TB results in the greatest bias, whereas the bias for TR+A case is very small. The bias for both TB and TR+A increases as effect sizes and censoring rates grow. Coverage of the true effect size for TB drops to zero for common variants (*MAF* = 0.4) already at *β*_1_ = 0.2. The coverage for TR+A is not as good as for TA, although the differences are minor. Coverage can be seen to be better consistently when MAF is low.

For power we only present the plot, where MAF=0.05 and CR=90 (Figure 2) as the differences in the results are greatest here, other plots can be found in the Supplement Figure 1.

**Figure 2.**
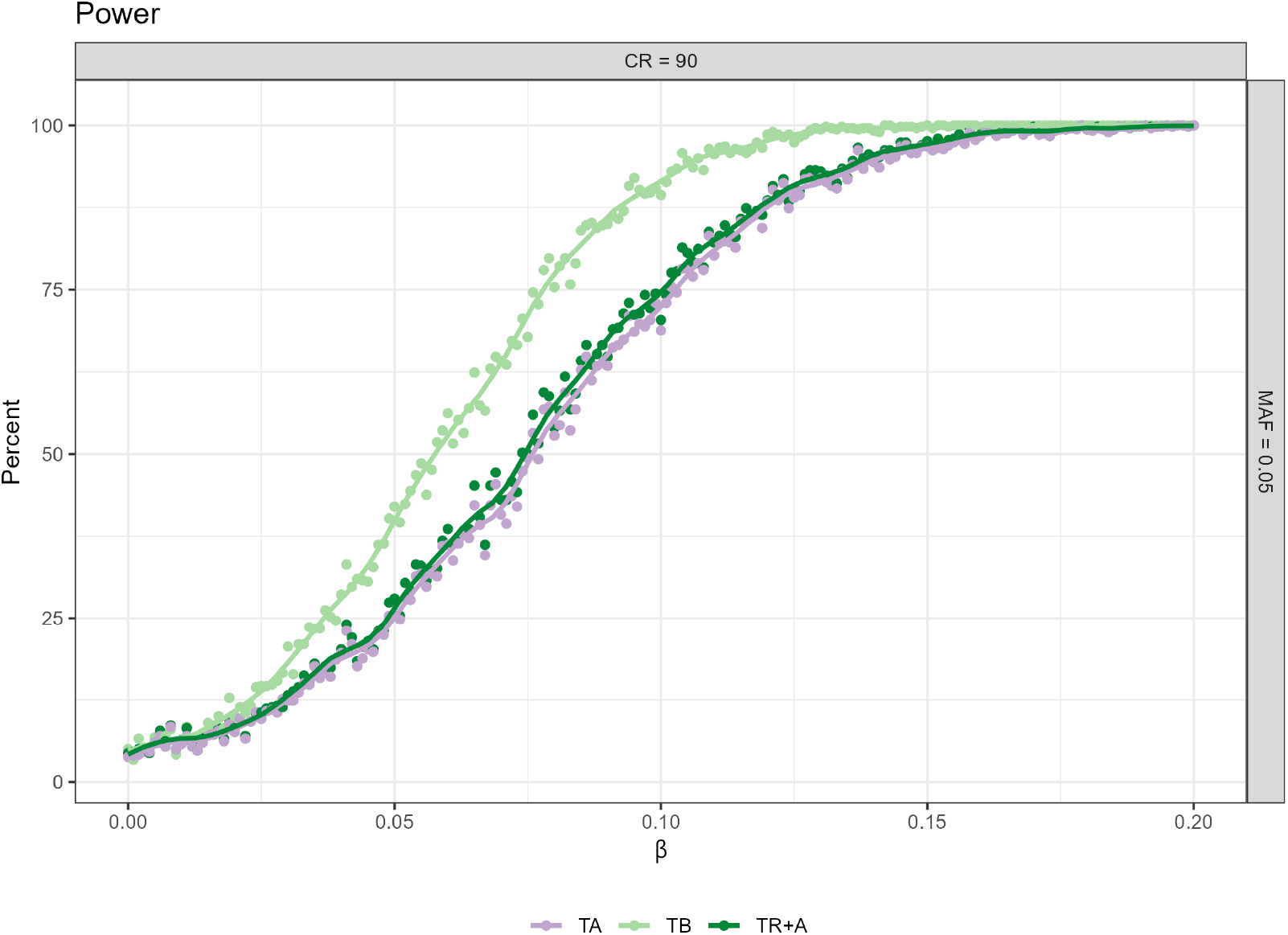
Power analysis for censoring rate (CR) 90% and minor allele frequency (MAF) 0.05 across three different timescale choices (TA, TB, TR+A). Additional power figures for varying censoring rates and MAFs are available in the Supplementary Figure 1.

CPH with TB results in the highest power to detect a significant association, whereas the power for TA is lowest no matter what the effect size. The differences in power for TB and TR+A can be up to 25%. CPH with TR+A and TA have very similar power regardless of the effect size.

As a conclusion we see that although the TB approach leads to potentially biased estimates of the true effect, it may be the preferred approach if the aim is to maximize power in a discovery study.

### 4.2 Utility of martingale residuals based approach in approximating CPH model estimates

As shown before, CPH with TB could be preferable in GWAS settings due to highest power to detect relatively small effect sizes. Therefore, the simulation results on the performance of the MR approach are here presented only for TB (Figure 3), whereas the results for CPH with TR+A and TA are presented in the Supplementary Figures 3 and 4.

**Figure 3.**
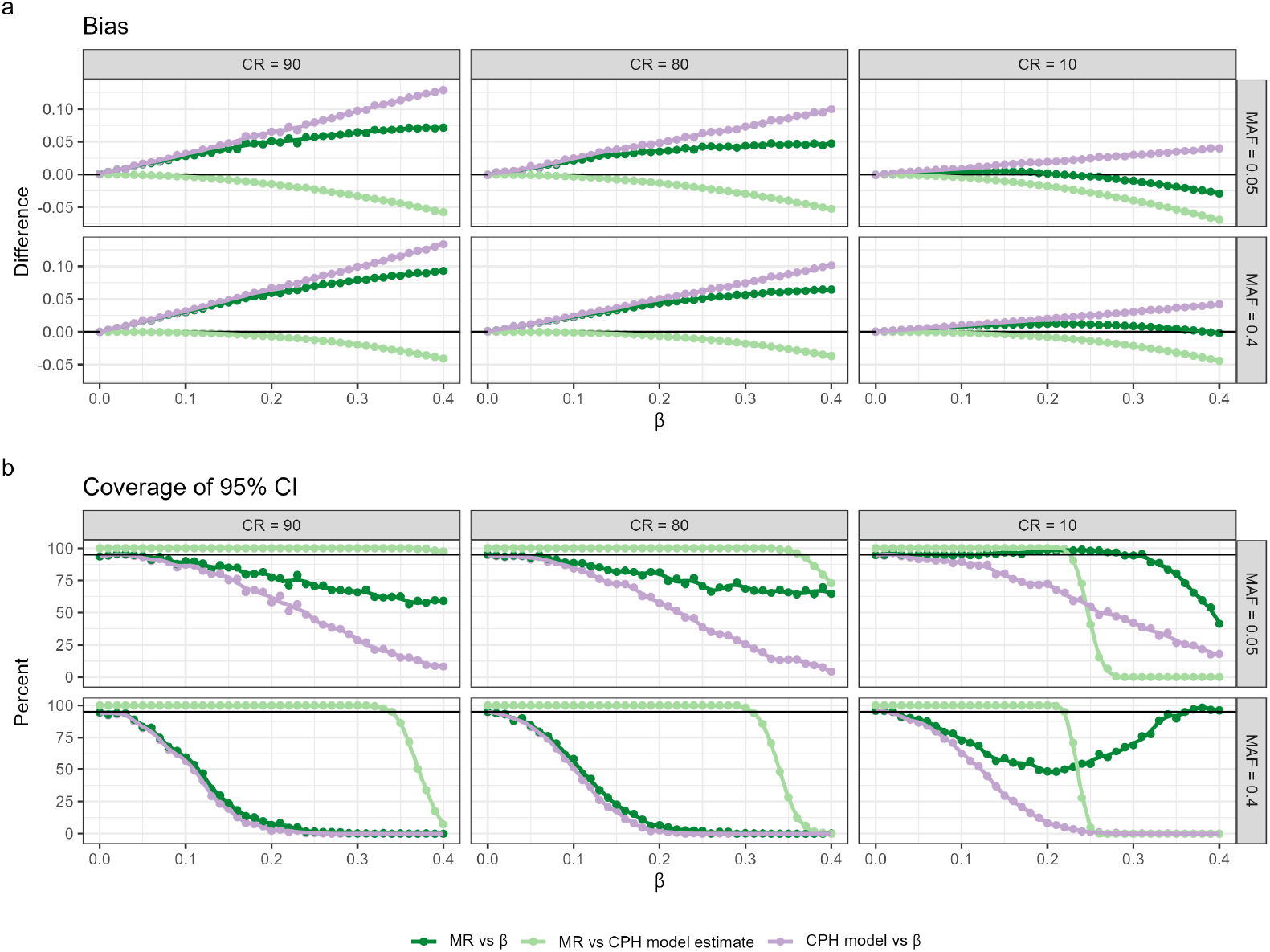
Bias and coverage of 95% CI for models using age since birth as the timescale (TB), evaluated across different censoring rates and MAFs. Three different scenarios are compared*β* estimate using MR approach vs true *β (MR vs β), β* estimate using MR approach vs CPH estimate *(MR vs CPH model estimate)* and CPH model estimate vs true *β (CPH model vs β)*. Results for other timescales are presented in Supplementary Figures 3 and 4.

A comparison of MR estimates with those from the standard CPH modelfitting algorithm and the actual effect sizes reveals that the two-step approach approximates CPH estimates quite well within typical GWAS working range. Compared to CPH estimates, the effect size estimates based on MR approach demonstrate less bias and higher relative accuracy in capturing the true parameter estimates. However,the relationship between bias and the true effect size exhibits a non-linear pattern, and the same nonlinearity holds for the coverage of the 95% CI. The observed nonlinearity requires further theoretical investigation. Similarly to the effect sizes, p-values obtained from the MR approach were more conservative than the ones obtained from the CPH model within typical GWAS setting. We observed in our simulation that when censoring rates were low, MR p-values approximated CPH p-values better, compared to settings with high censoring rates. Nevertheless, the power is rather similar to the CPH across different censoring rates and MAF for effect sizes within GWAS working range (see Supplementary Figure 2).

### 4.3 Relatedness and model misspecifation

To investigate issues concering relatedness and model misspecification, we simulate datasets of siblings with three covariates to mimic realistic genotype, phenotype, and a shared family frailty, which is often unmeasured. The simulation details are provided in the Supplement Section 2.2. We run analyses using three different timescales (TB, TR+A, TA), both ignoring relatedness (ie including relatives) and using only unrelated individuals. For each scenario, we compare models including all covariates to those omitting the frailty term. Additionally, we evaluate a CPH model with a frailty term for data including relatives, but ignoring frailty term as it would be in a realistic setting. Results indicate that ignoring relatedness does not significantly increase bias in effect size compared to the choice of timescale, regardless of censoring rate or MAF. However, omitting a covariate creates substantial bias and, in our setting, even changes the direction of the bias. The frailty CPH model shows the smallest bias when using an age-adjusted timescale, with sensitivity to censoring rate and MAF—the smaller these two, the smaller the bias. Power analyses reveals the highest power for large censoring and small MAF occurres with a birth-based timescale. Power is sensitive to censoring rate, MAF, ignoring relatedness, and omitting covariates, with the greatest decrease observed when both relatedness were ignored and covariates omitted. Notably, the coverage of the 95% CI decreases sharply when a covariate is omitted and relatives are included in the analysis, particularly when left-truncation is accounted for. This effect is more pronounced if the censoring rate decreases and MAF increases, due to the underestimation of variance (see Supplementary Figure 5).

#### Martingale residuals in relatedness and model misspecification

We investigate how the two-step martingale residual approach would work in a realistic setting by using a simulation in which we intentionally omit a covariate representing frailty. We are interested in whether the two-step martingale residual approach could approximate CPH estimates, specifically using age since birth as the timescale and under high censoring conditions. This investigation is conducted for both related and unrelated subjects. For this setup, the martingale residual approach leads to the smallest bias and highest coverage, although it has slightly less power than the standard CPH models (Figure 5). Therefore, for explanatory GWAS using age since birth as the timescale, the martingale residual-based approach appears to be a robust method for estimating hazard ratios, effectively handling related subjects and being computationally efficient.

**Figure 4.**
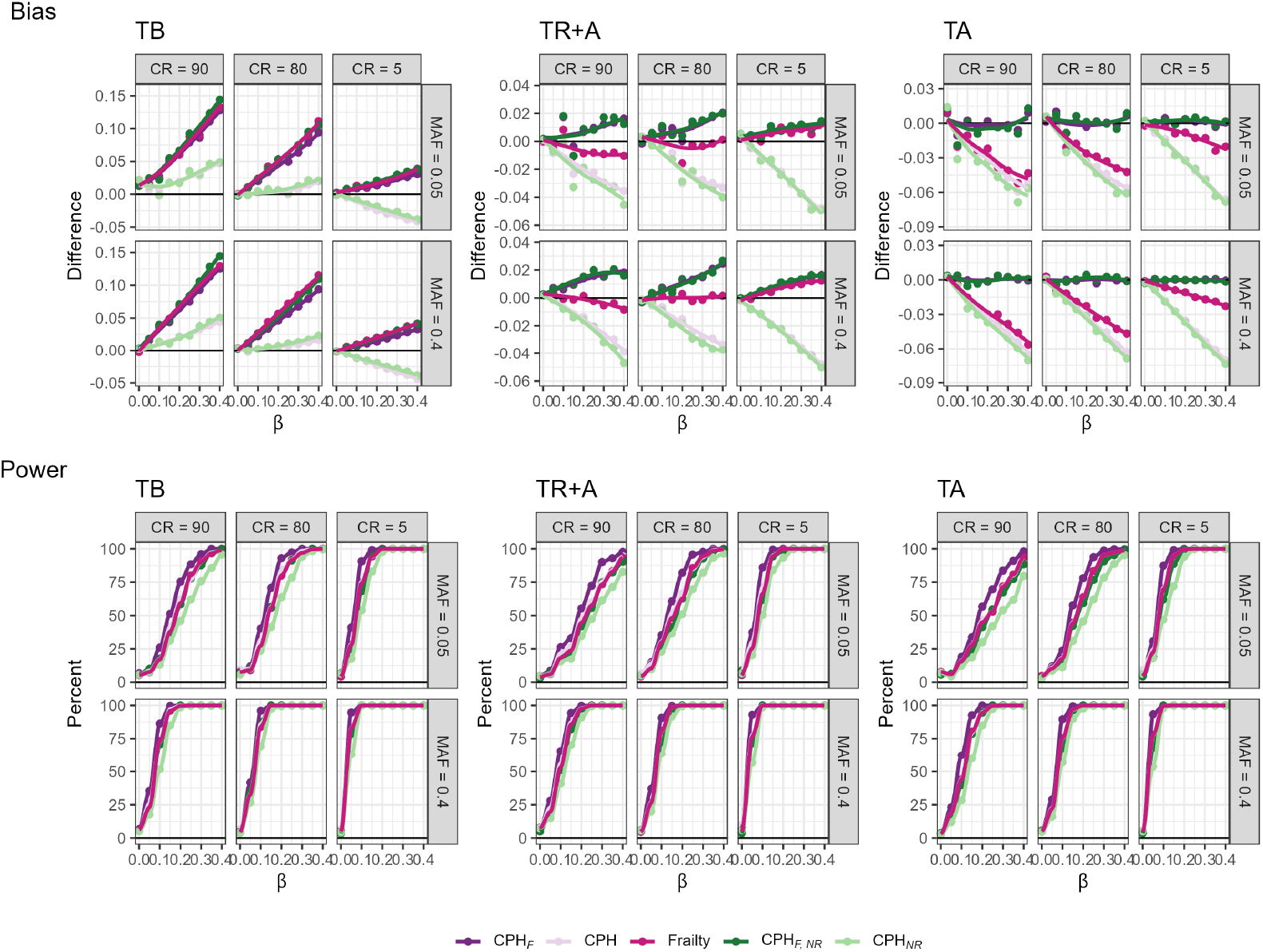
Bias and power of various models across three timescales (TB, TR+A, TA), different censoring rates (CR), and MAFs fitted on a cofort with related individuals. The models include full covariate models with all subjects (*CPH*_*F*_) and only independent subjects (*CPH*_*F*,*NR*_), models omitting the frailty term for all subjects (*CPH*) and only independent subjects (*CPH*_*NR*_), and a CPH frailty model for all subjects (*Frailty*).

**Figure 5.**
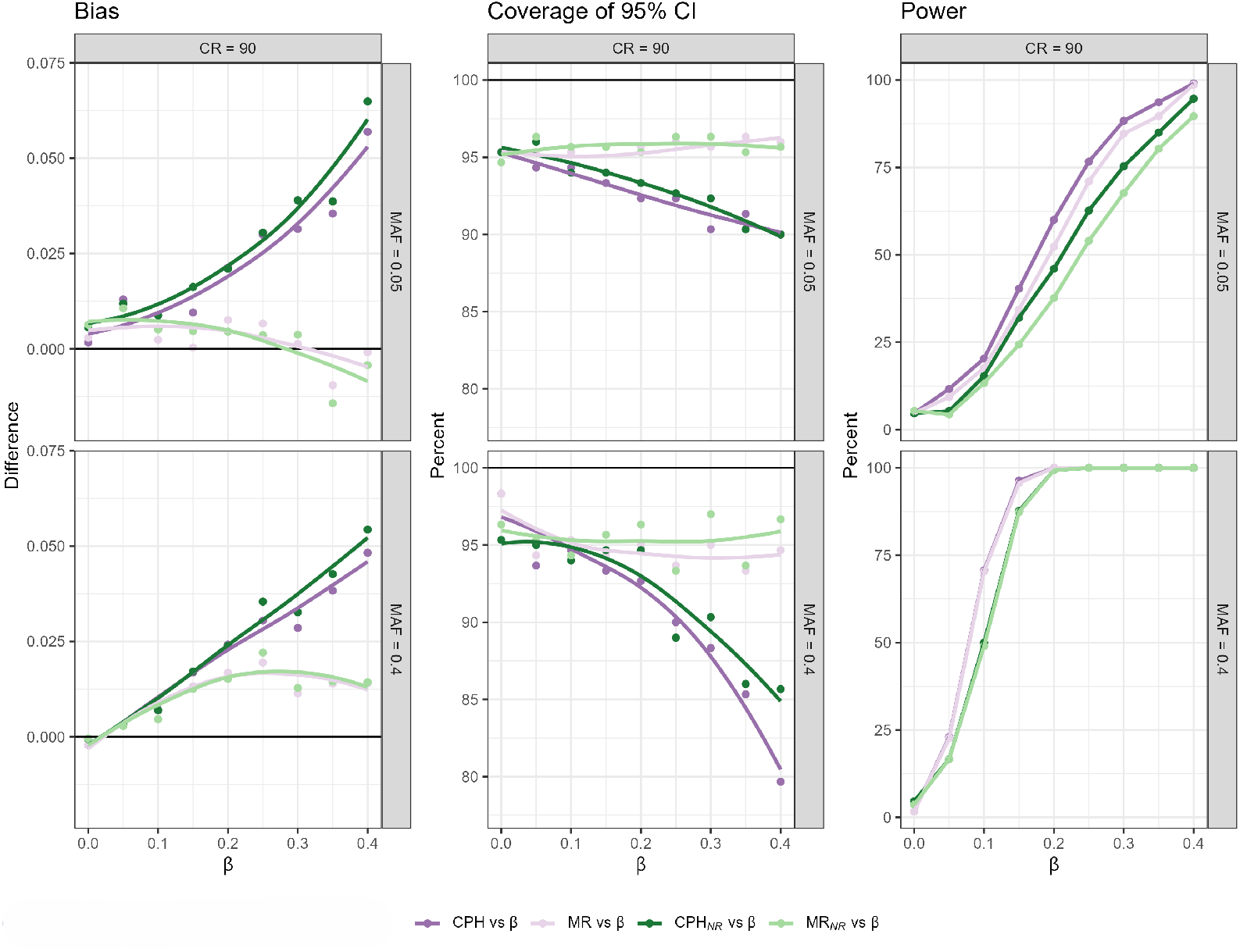
Bias, coverage of 95%CI and power, resulting from fitted models that use age since birth as a timescale, across different censoring rates (CR), and MAFs in a cohort including related individuals. Four different scenarios are comparedCPH model and MR-based model fitted on the complete dataset *(CPH vs β, MR vs β)*, and respective models fitted on the subset with independent subjects (*CPH*_*NR*_ *vs β, MR*_*NR*_ *vs β*).

### 4.4 Application to the Estonian Biobank data

The Estonian Biobank maintains a volunteer-based cohort of the Estonian adult population (aged ≥ 18 years) [40]. The sample size used in this analysis is 51 463, which represents approximately 5% of the Estonian adult population (participants recruited during the first period of recruitment in 2003-2011). In this sample, 65.6% of participants were female and the median age at recruitment was 43 (min = 18, max = 103) years. Median follow-up time with IQR was 13.1 (11.7; 13.9) years. The lifespan data of the participants is obtained via record linkages with the Estonian Causes of Death Register (latest linkage for the data used here was in the beginning of 2022). The mortality rate in the analysed sample was 13.2%.

Testing the top three SNPs and the polygenic risk score (GRS) for lifespan based on Timmers et al. [23], we fit models with three choices for timescale: time since birth (TB), time since recruitment with age-adjustment (TR+A) and age as timescale (TA, accounting for left-truncation). For each timescale choice, we fit the model for the entire sample (ignoring the relatedness) and also for the sample where relatives are excluded (the remaining sample size: *n* = 38 223). Thus as a result, 6 different models are fitted in total.

Out of the 3 top SNPs tested, only one (APOE) shows significant result in the Estonian Biobank (Figure 6). As expected the GRS shows the greatest effect on mortality, whereas its effect size is almost identical regardless of whether the relatives are included or excluded. Largest effect size estimate is obtained when TB was used as the timescale – as pointed out before, the bias due to left-truncation is a likely cause for this difference. The other two timescale choices do not lead to visible differences in the effect sizes. The estimates of the effects of the LPA, APOE and CHRNA3/5 variants do not differ much, but excluding relatives has generally reduced the estimated effect sizes for APOE and CHRNA3/5, whereas the timescale choice does not really have any clear impact.

**Figure 6.**
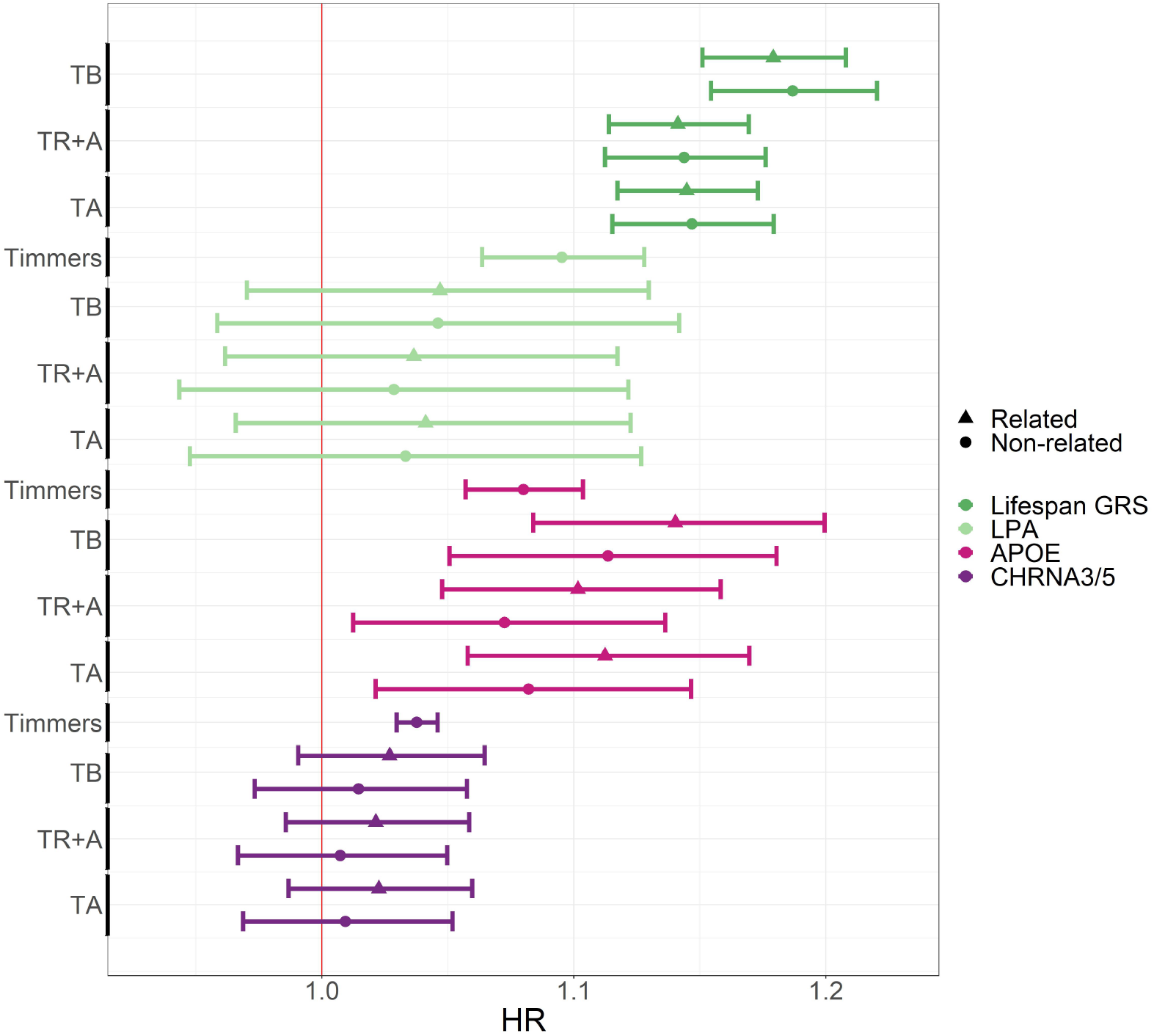
Comparison of hazard ratios for survival-associated SNPs and lifespan GRS using CPH with different timescales and related vs unrelated participants in the Estonian Biobank data. The figure shows HR estimates with 95% confidence intervals x-axis, and the timescale choice or eLife article reference on the y-axis. Gene names, rather than SNP names, are indicated by color and relatedness by shape. TB, TR+A and TA correspond to CPH using time since birth as timescale, time since recruitment + age adjustment as timescale and age as timescale with all EstBB participants.

## 5 Discussion

For accurate estimation of population parameters, unbiasedness is the essential requirement for any statistical estimators. However, the task of estimation should be distinguished from the task of hypothesis testing. The present work has highlighted that in the context of GWAS for time to event outcomes, the estimators leading to the smallest bias are not necessarily the ones corresponding to most powerful tests for the hyopthesis of no genotypephenotype association.

Time-to-event phenotypes are challenging for GWAS, as the commonly used analysis tools, such as CPH modeling, require considerably more computational resources for implementation than algorithms for linear and logistic regression analysis. In addition, as the power depends not on the total sample size, but on the number of (disease or mortality) events observed, even a large biobank cohort may not be sufficiently powered for the discovery of biologically meaningful outcome-associated variants. Thus approaches that maximize power are especially welcome for time-to-event GWAS analyses, even if they come at the cost of some bias in parameter estimates.

The first finding of the present study is, that although careful adjustment for left-truncation is needed to achieve unbiasedness, it would considerably reduce power in a discovery GWAS compared to the approach that ignores it. For instance, a true hazard ratio of 1.05 (typical effect size of a common variant in GWAS) is likely to be overestimated by 2-3%, whereas the power to detect an association may be increased by more than 1.5 times, when time since birth is used as timescale and left-truncation is ignored.

As under the null hypothesis of zero effect size the bias could not occur, ignoring left-truncation would not increase type I error probability. There is, however, one exception – the case where a genetic variant has been under selection. By “enriching” the risk sets with individuals recruited at later time points, one may in these cases create a situation where the allele frequencies in subjects with outcome events differ systematically from the allele frequencies in the risk sets. We recommend that this issue should be examined for variants identified as significant in a GWAS.

To simplify the computational algorithm of model-fitting, the use of a twostep procedure involving martingale residuals has been explored in the GWAS context. Martingale residuals were initially proposed as a diagnostic tool for a CPH model (mainly to identify appropriate covariate transformations), and to our knowledge, their use in the actual effect estimation has not been explored in detail. We have shown that the two-step procedure provides valid estimates with no (or negligible) bias for the estimation of the relatively small effect sizes that are typical for GWAS findings. In addition, we have also shown that using the MR approach the power of the association discovery is not decreased compared to the corresponding CPH model.

Based on our simulations, ignoring relatedness does not significantly increase bias in effect size, whereas omitting key covariates introduces substantial bias. Additionally, the two-step martingale residual approach proved to be robust for estimating hazard ratios, efficiently handling related subjects with high coverage and slightly reduced power compared to standard CPH models.

In summary, our results support that for a time-to-event phenotype, a procedure where: 1) age is used as a time-scale and left-truncation is ignored and 2) a two-step procedure that obtains martingale residuals at the first step and runs a linear regression-based GWAS as the second step is implemented, leads to better computational efficiency and better power for variant discovery than the procedure that fits a CPH model separately for each variant, whereas adjusting for other covariates.

Once the set of potentially associated variants is identified, we still recommend to validate the findings in both the discovery cohort(s) and also in a large independent cohort, using the CPH modeling approach that leads to unbiased estimates (thus, properly accounting for left-truncation). The latter is especially true, when the effects of polygenic risk scores (GRS) are estimated, as biases in these estimates are not acceptable when personalized risk prediction algorithms are derived. Also, to compute a GRS based on estimated regression coefficients from GWAS, one needs unbiased estimates for those coefficients.

## Acknowledgements

Anastassia Kolde was supported by funding from the European Union’s Horizon 2020 research and innovation programme under the Marie Sklodowska-Curie grant agreement No. 813533. Krista Fischer, Merli Koitmäe and Meelis Käärik received support from Estonian Research Council through grant PRG1197. Merli Koitmäe recieved support from Estonian Research Council through grant PRG1255. This work was partially written at writing retreats and writing days organised by The University of Tartu Institute of Genomics

